# Transcriptional condensates at super-enhancers mediate pH-dependent transcriptional control in innate immunity

**DOI:** 10.1101/2025.10.29.685293

**Authors:** Shengyuan Wang, Zhongyang Wu, Zhe Zhong, Xu Zhou, Chongzhi Zang

## Abstract

Tissue acidification is a common feature of hypoxia, inflammation and solid tumor. Acidic pH regulates innate immune response in macrophages by weakening BRD4-containing transcriptional condensates. Yet how disruption of transcriptional condensates leads to gene-specific regulation of immune programs remain unclear. Here, we integrated ATAC-seq, ChIP-seq, and RNA-seq of primary murine macrophages and performed integrative epigenomics analyses to identify transcriptional regulators (TRs) with pH-sensitive regulatory potential and association to BRD4-dependent transcriptional condensates. We determined pH-dependent super-enhancers (SEs) by extended profiles of BRD4 binding and h3K27ac marks. We found RELA, IRF family, and STAT family as candidate TRs enriched at BRD4-associated, pH-sensitive SE regions. RELA and IRF3 preferentially occupied BRD4-associated and pH-sensitive SEs, and displayed markedly reduced binding under acidic conditions, aligning with BRD4 occupancy change. Correspondingly, immune-response genes within BRD4-associated, pH-sensitive SE regions, including *Ch25h*, *Acp2*, *Gda*, and *Ifit* family, were significantly higher expressed at pH 7.4 than at pH 6.5. Together, these results reveal a set of TRs involved in BRD4-associated, pH-sensitive transcriptional condensates that coordinate macrophage gene activation under physiological conditions, providing mechanistic insight into how acidic stress modulates transcriptional condensates and immune responses.

## Introduction

Gene regulation is fundamental to cellular function, enabling cells to respond dynamically to developmental cues, environmental signals, and pathological conditions. Precise control of gene expression programs relies on coordinated regulation between distal regulatory elements and their target genes, mediated by complex assemblies of transcriptional machinery. In particular, enhancers are critical *cis*-regulatory elements that control spatiotemporal gene expression patterns often from considerable genomic distances. Recent advances in understanding enhancer function have revealed that enhancers do not operate in isolation but rather function through dynamic three-dimensional chromatin interactions.

Among enhancers, super-enhancers (SEs) represent a specialized class of regulatory elements that play particularly important roles in controlling cell identity and stimulus-responsive gene expression programs. They are clusters of enhancers that span broad genomic regions and are characterized by high H3K27ac and high occupancy of acetyl-lysine readers such as BRD4 and BRD9^1–3^. SEs contain many cis-regulatory elements bound by multiple transcription factors (TFs), which recruit the general transcription machinery and coactivators to drive robust expression of target genes^4^. One emerging mechanism underlying the heightened transcriptional activity at super-enhancers involves liquid-liquid phase separation (LLPS), whereby transcriptional condensates, the dense, dynamic assemblies of transcriptional regulators (TRs), including TFs and chromatin regulators, could form at these genomic loci^5–8^. These biomolecular condensates represent a distinct mode of cellular organization that compartmentalizes and concentrates key regulatory factors to facilitate efficient gene activation. Many condensate components harbor intrinsically disordered regions (IDRs) that enable multivalent, weak interactions necessary for condensate formation^9^. These condensates create specialized microenvironments that promote robust transcriptional activation by concentrating RNA polymerase II, Mediator complex, and other coactivators at active genes.

Despite these advances, critical questions remain regarding which TRs participate in these condensates and how their assembly and function are regulated in response to cellular and environmental signals. Our recent work^10^ as identified intracellular pH as a novel regulator of transcriptional condensates in macrophages. Specifically, the bromodomain-containing protein BRD4 acts as a pH-sensitive regulator through its histidine-enriched intrinsically disordered region. At physiological pH (7.4), BRD4 forms stable condensates with MED1 and other TRs at SEs marked by broad H3K27ac. However, acidic conditions (pH 6.5) disrupt these multivalent interactions, leading to dissolution of BRD4-MED1 transcriptional condensates. Consistent with this pH-dependent mechanism, BRD4 occupancy at key immune-regulatory loci with broad H3K27ac patterns is markedly reduced at pH 6.5 compared to pH 7.4, suggesting that pH regulates condensate-dependent transcriptional activation at super-enhancers.

In this study, we performed integrative epigenomic analyses to identify TRs related to BRD4-associated, pH-sensitive transcriptional condensates in primary murine macrophages. Integrating ChIP-seq, ATAC-seq, and transcriptomic data, we identified RELA, and IRF and STAT families as candidate pH-sensitive transcriptional regulators. Using ChIP-seq profiled at pH 7.4 and pH 6.5, we found that RELA and IRF3 exhibit significantly reduced binding at BRD4-associated, pH-sensitive SEs under acidic conditions. These findings support a model in which multiple transcriptional regulators contribute to BRD4-associated condensate function, revealing a coordinated mechanism by which environmental pH modulates inflammatory gene expression programs through condensate dynamics and transcriptional network.

## Results

BRD4 plays a critical role in regulating immune responses through the formation of transcriptional condensates. Under acidic conditions, BRD4 activity is disrupted, leading to the dissolution of BRD4-associated condensates and altered target gene expression (Figure 1A). To uncover the genomic basis of this regulation, we identified differential BRD4 binding sites at super-enhancer (SE) regions between the two pH conditions using region-level differential analysis (Figure 1B). Because BRD4 functions as a reader of acetylated histones, many differential sites may reflect changes in histone acetylation rather than BRD4-specific regulation. To focus on BRD4-dependent effects, we narrowed our analysis to sites with non-differential H3K27ac signals. We considered transcriptional regulators (TRs) showing significant enrichment at these sites as those associated with BRD4 condensates. Through multiple integrative epigenomic analyses, we identified several TRs that are functionally linked to BRD4 condensate regulation.

**Figure 1.**
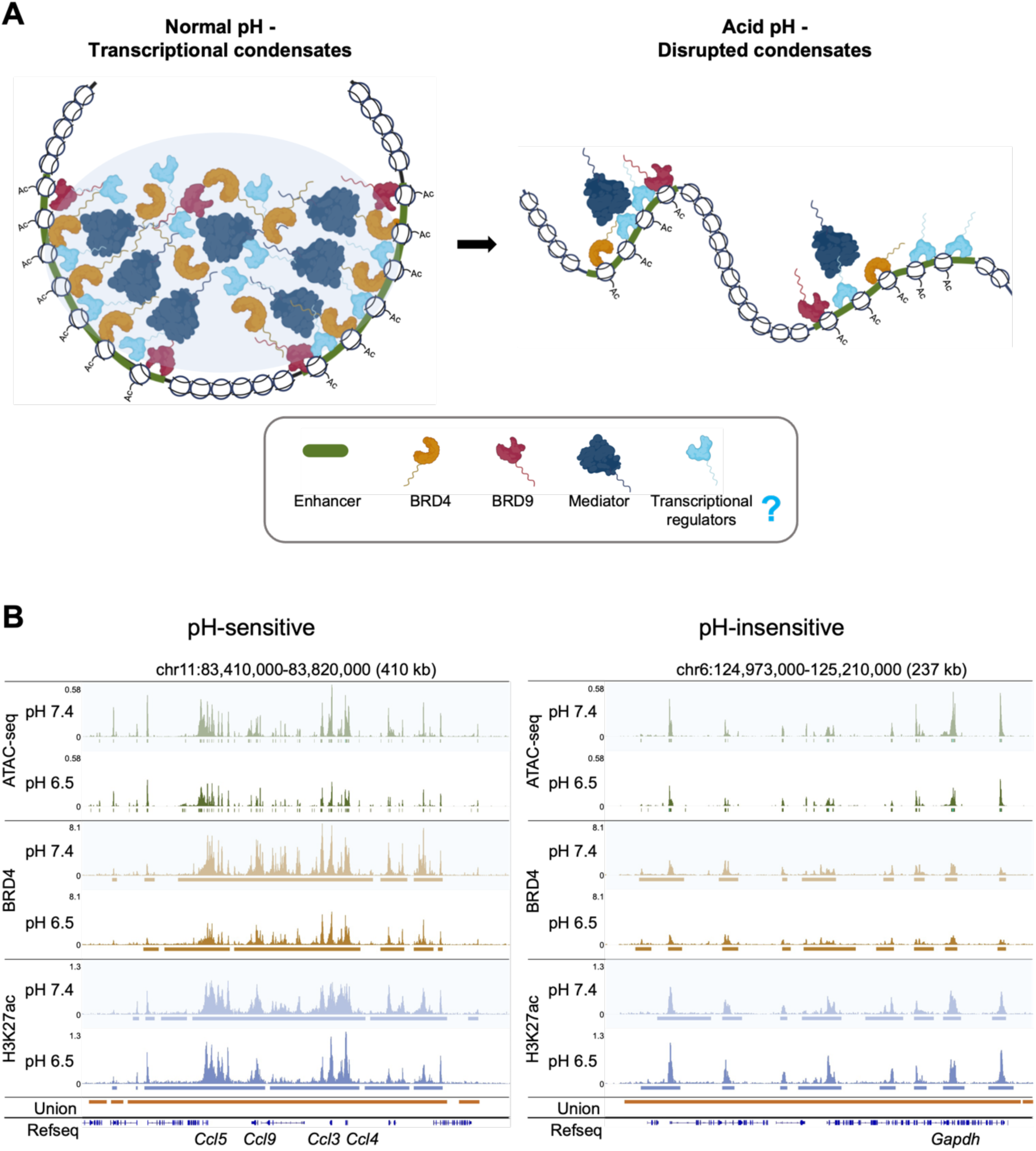
BRD4-associated, pH-sensitive transcriptional condensates at super-enhancer (SE) sites. (**A**) Schematic illustration of BRD4-associated, pH-sensitive transcriptional condensates at SE regions. Under physiological pH (7.4), transcriptional condensates form at SE sites characterized by high histone acetylation, increased chromatin accessibility, and recruitment of transcription factors, BRD4, BRD9, and Mediator complexes, resulting in high local molecular concentration and condensate formation through liquid–liquid phase separation. Under acidic pH (6.5), histone acetylation remains unaffected, but BRD4 is dissolved, disrupting condensate integrity. The goal of this study is to identify transcriptional regulators (TRs) that participate in BRD4-associated, pH-sensitive transcriptional condensates. (**B**) ChIP-seq tracks of chromatin marks across 2 pH conditions at union SE sites. The left panel shows a pH-sensitive site at the *Ccl* locus, where BRD4 binding and Ccl genes expression are significantly higher at pH 7.4 than at 6.5, while the right panel shows a pH-insensitive site at the housekeeping gene *Gapdh* locus, where BRD4 binding and gene expression is unchanged across pH. In both cases, H3K27ac levels remain stable.

### Region-centric analysis identified multiple TRs related to BRD4-associated, pH-sensitive transcriptional condensates

To identify TRs related to BRD4-associated, pH-sensitive transcriptional condensates, we first defined putative SE sites in the mm10 mouse genome. As SEs represent clusters of enhancers characterized by high histone acetylation and chromatin accessibility, we generated union sites by merging peaks from three chromatin marks, including ATAC-seq, BRD4, and H3K27ac across all experimental conditions, and considered the union sites with at least 10 kb in length as putative SEs, resulting in 15,061 sites (median length: 19 kb; range: 10–556 kb).

To characterize BRD4-associated transcriptional condensates that are disrupted in acidic condition, we sought to identify differential BRD4 occupancy that is significantly higher at pH 7.4 compared to pH 6.5 (Figure 2A). Using SICER^11^ for differential analysis, we identified SEs with significantly higher BRD4 signals under neutral pH (Figure 2B). To specifically characterize pH-sensitive transcriptional condensates associated with BRD4 but not general H3K27ac defined SEs, we defined the SE sites showing significantly higher BRD4 but not higher H3K27ac levels at pH 7.4 than pH 6.5 as pH-sensitive condensate-associated SEs, and sites without significant enrichment for either BRD4 or H3K27ac as pH-insensitive SEs. We generated pH-sensitive and pH-insensitive SE sets separately for LPS-activated and control samples, yielding 514 pH-sensitive and 5,815 pH-insensitive SEs under LPS stimulation, and 287 pH-sensitive and 6,169 pH-insensitive SEs under control conditions.

**Figure 2.**
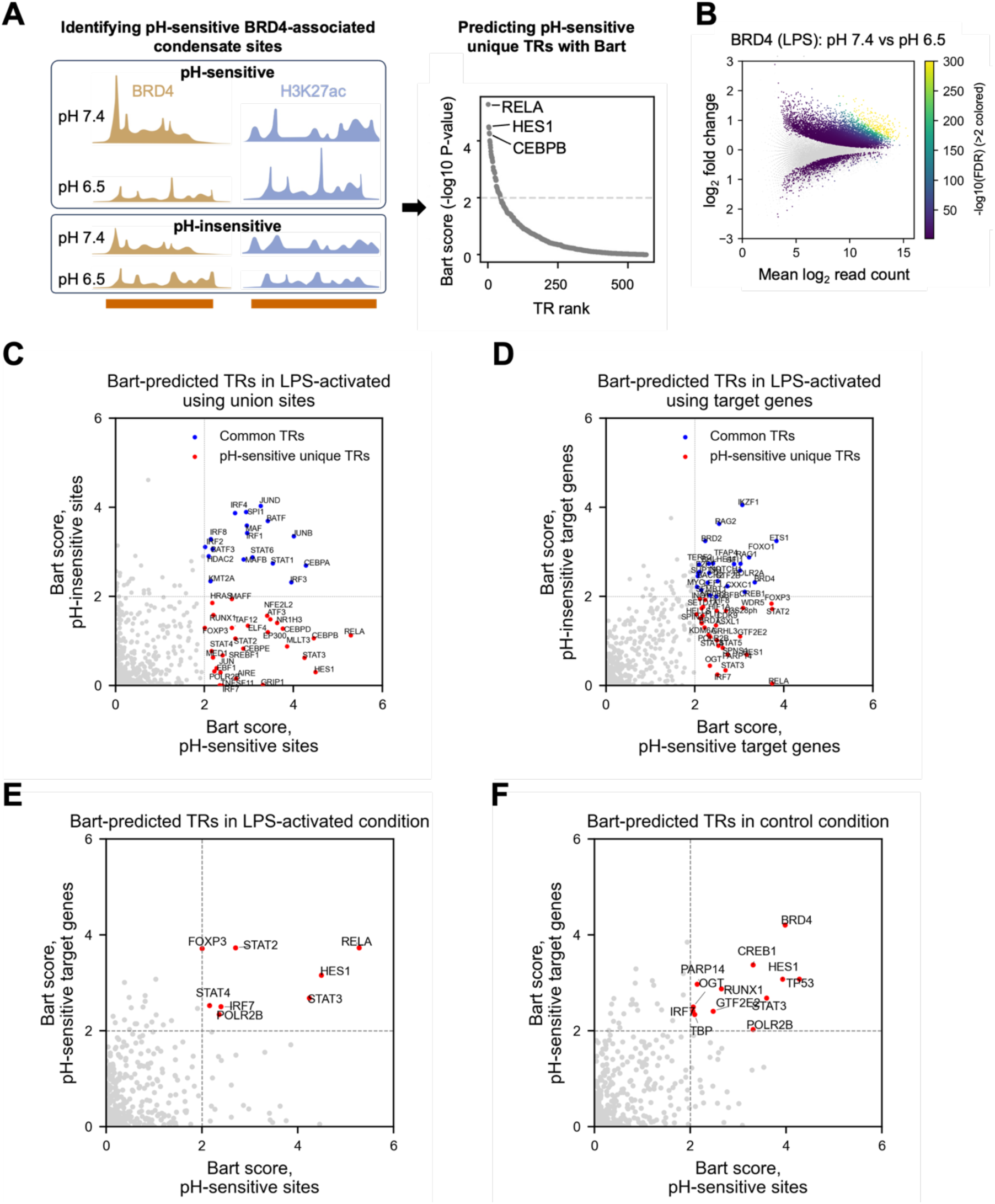
Region-centric analysis identifies TRs associated with BRD4-dependent, pH-sensitive transcriptional condensates. (**A**) Workflow of the region-centric analysis. pH-sensitive regions were defined as SE sites with significantly higher BRD4 signal at pH 7.4 than 6.5 but without significant H3K27ac change, while pH-insensitive regions showed no significant changes in either mark. BRD4-associated TRs were predicted using BART based on pH-sensitive-specific enrichment. (**B**) Differential analysis using SICER2 identified BRD4 and H3K27ac differential SE sites across pH conditions. MA plot shows SE sites with average normalized read counts versus log₂ fold changes; significantly enriched sites are color-coded by FDR. (**C**, **D**) BART analysis identified TRs associated with pH-sensitive and pH-insensitive regions (**C**) and their target genes (**D**) under LPS activation. (**E**, **F**) TRs uniquely enriched in pH-sensitive regions and their target genes under LPS stimulation (**E**) and control (**F**) conditions.

We then identified TRs potentially bound at these pH-sensitive and pH-insensitive SEs by applying BART^12^ with these SEs as region set input (Figure 2A). We considered the identified TRs with P-value < 0.01 and uniquely enriched in pH-sensitive SEs as BRD4-condensate– associated TRs, and those shared by both pH-sensitive and pH-insensitive SEs as general SE-associated TRs (Figure 2C).

Next, we identified putative target genes for both pH-sensitive and pH-insensitive SEs by calculating regulatory potential (RP) scores^13^ for genes linked to each SE set, defined as the distance-weighted sum of normalized ChIP-seq signals. We predicted TRs associated with these target genes using BART (Figure 2D) and regarded TRs identified by both region-and gene-based analyses as high-confidence regulators of Brd4-associated, pH-sensitive transcriptional condensate (Figures 2E, F).

Applying this approach to the datasets at LPS stimulation, we identified eight TRs, including RELA, IRF7, HES1, FOXP3, STAT family (STAT2, STAT3, STAT4), and POLR2B (Figure 2E). Among these, HES1 and FOXP3 are not expressed in macrophages and are disregarded.

RELA has previously been reported to show reduced binding at distal enhancer regions under acidic conditions, consistent with condensate dissolution. STAT family members have critical functions in immune regulation^14^. Under control conditions (without LPS stimulation), we identified IRF7, STAT3, TP53, and POLR2B as major regulators (Figure 2F). Notably, TP53 is known to participate in transcriptional condensates and to play important roles in immune and stress responses^15,16^. We further examined whether the identified TRs possess intrinsically disordered regions (IDRs) using AIUPred^17,18^ and found that all contained IDRs (Supplementary Figure S1), suggesting their potential to form transcriptional condensates.

Together, we identified multiple TRs involved in immune regulation and potentially engaged in BRD4-associated, pH-sensitive transcriptional condensate formation through region-centric analysis.

### Consistent identification of TRs at BRD4-associated, pH-sensitive SEs across multiple analyses

Compared with the region-centric analysis, which begins by identifying genomic regions exhibiting BRD4-associated condensate patterns, the gene-centric analysis focuses on transcriptional regulation from the perspective of each gene. We calculated the RP for each gene, and considered the genes with significantly higher RP for BRD4 under pH 7.4 than pH 6.5, but not for H3K27ac as target genes that are regulated by BRD4-associated transcriptional condensates (Figure 3A). We then identified the TRs that regulate these genes using BART. We identified RELA and STAT3 as key TRs related to BRD4 condensates (Figure 3B), consistent with results from the region-centric approach. Other TRs, including IRF7, STAT2, and POLR2B, which were detected in the region-centric analysis, did not meet our stringent cutoff of P-value < 0.01 in gene-centric analysis but still had the P-value < 0.05, indicating the consistency.

**Figure 3.**
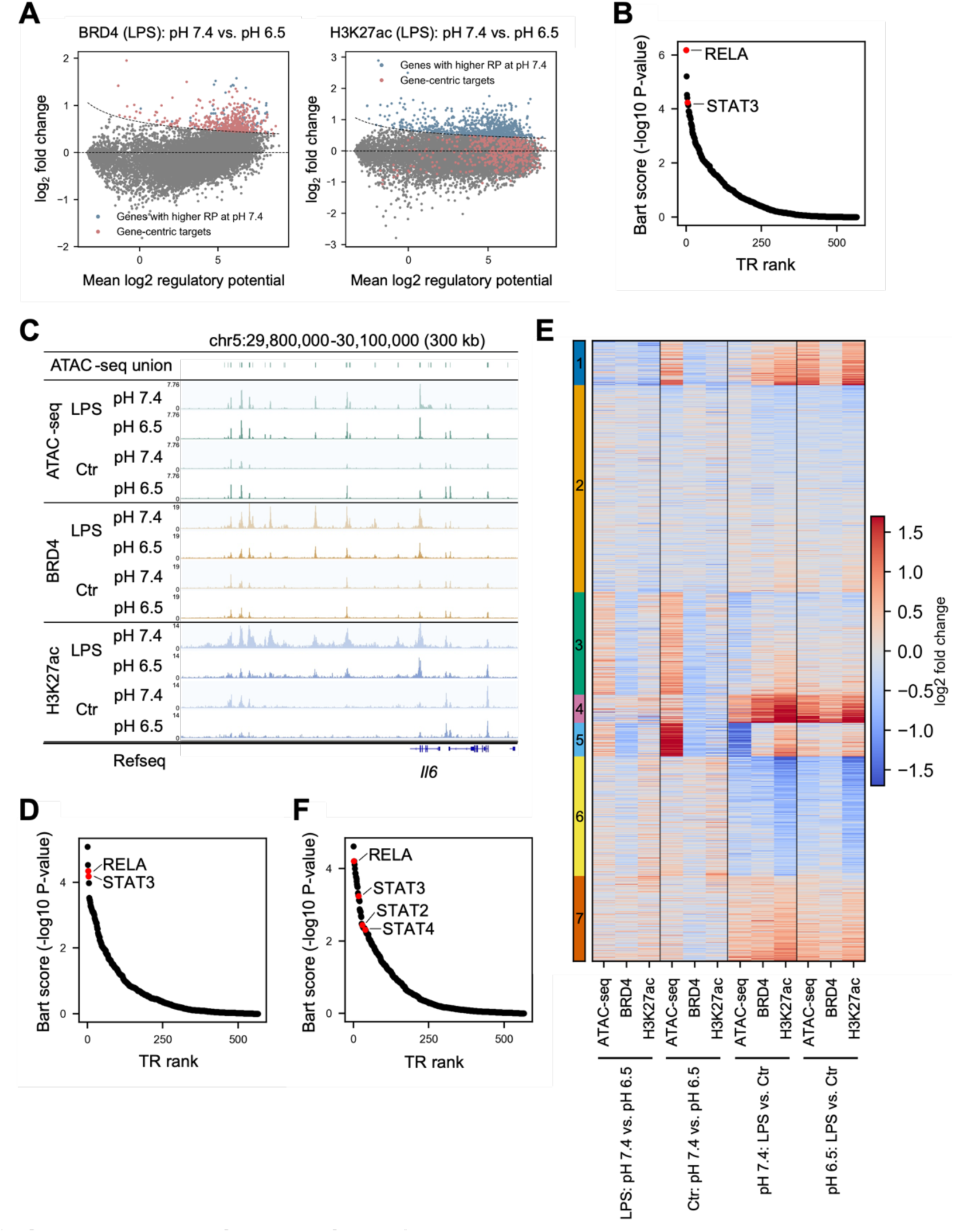
Consistent identification of BRD4-associated, pH-sensitive TRs across complementary analytical frameworks. (**A**) Gene-centric analysis identified BRD4-associated, pH-sensitive target genes showing significantly higher regulatory potential (RP) scores for BRD4 at pH 7.4 but not for H3K27ac. Pink dots mark genes meeting the BRD4-specific criteria, blue dots represent genes showing significant differences for either mark, which beyond the theoretical curve. (**B**) TRs predicted by gene-centric analysis, with highlighted factors also identified in the region-centric analysis. (**C**) Illustration of the open chromatin-centric analysis using *Il6* locus as an example. ATAC-seq union sites were generated by merging ATAC-seq peaks across all conditions. (**D**) TRs identified by the open chromatin-centric analysis, with highlighted factors overlapping those from the region-centric analysis. (**E**) Unsupervised clustering grouped SE sites into seven clusters based on the log₂ fold changes of ATAC-seq, BRD4, and H3K27ac signals across conditions. (**F**) TRs enriched in cluster 4 ATAC-seq union peaks, with highlighted factors shared with the region-centric analysis.

To identify TRs based on their potential genomic binding sites, we performed an open chromatin-centric analysis focusing on accessible chromatin regions. We generated ATAC-seq union sites by merging all ATAC-seq peaks across conditions, and normalized read counts for each union site within each condition (Figure 3C). Because BRD4 and H3K27ac typically occupy broader genomic domains, we considered the BRD4 and H3K27ac signals in a broad region ±2 kb from the center of each ATAC-seq union site. Following a similar strategy as the region-centric analysis, we defined the sites showing significantly higher BRD4 signal at pH 7.4 than pH 6.5 but not higher H3K27ac signal as pH-sensitive open chromatin regions, and the sites without significant enrichment for either mark as pH-insensitive open-chromatin regions, then we identified target genes by calculating RP and predicted TRs uniquely associated with both pH-sensitive regions and their target genes by BART.

In the open chromatin-centric analysis, we identified RELA and STAT3 as major TRs (Figure 3D), consistent with the results from the region-centric analysis. Additional TRs including IRF7, POLR2B, STAT2, and STAT4 were detected only in the region-centric analysis, although they remained significant (P-value < 0.05) in the open chromatin-centric results. Conversely, several TRs were uniquely identified by the open chromatin–centric approach, including CEBPA, IRF3, and other STAT family members. Many of these TRs were also present within region-centric pH-sensitive regions and are functionally linked to immune responses.

To further explore the overall chromatin landscape without imposing constraints from BRD4 or H3K27ac differential analyses, we examined the landscape of all broad regulatory regions as candidate SEs. For each SE, we calculated the log2 fold changes of ATAC-seq, BRD4, and H3K27ac signals between the two pH conditions and between LPS-stimulated and control samples to assess the effects of pH and LPS stimulation. We then performed unsupervised clustering using k-means (k = 7) to classify the regions into seven distinct clusters based on their log2 fold-change patterns (Supplementary Figure S2A). Each cluster exhibited unique chromatin patterns (Figure 3E). For instance, Clusters 4 and 7 showed consistent increase of all three chromatin marks under LPS stimulation compared with control, and BRD4 signals were higher at pH 7.4 than at pH 6.5 under LPS-stimulated conditions. In contrast, Clusters 3 and 5 displayed lower BRD4 signals at pH 7.4 compared with pH 6.5 under both LPS and control conditions. Using the pH-sensitive and pH-insensitive SE regions defined in the region-centric analysis, we performed enrichment analysis to determine their distribution across clusters. As expected, pH-sensitive SE regions were significantly enriched in Clusters 4 and 7, whereas pH-insensitive SE regions were enriched in Clusters 1, 3, and 5 (Supplementary Figure S2B), consistent with the observed dynamic chromatin patterns.

Since Clusters 4 and 7 were enriched for pH-sensitive sites, we used all ATAC-seq union sites located within SE regions in these clusters as input for BART in region mode to predict enriched TRs (Figure 3F; Supplementary Figure S2C). Most TRs identified through the region-centric analysis were also recovered by this unsupervised clustering approach, whereas pH-insensitive SE regions enriched Cluster 3 and 5 did not yield similar TRs, further supporting the robustness and specificity of the identified BRD4-associated, pH-sensitive TRs.

In summary, we performed region-centric, gene-centric, and open chromatin-centric analyses, each used distinct strategies to identify TRs associated with BRD4-defined, pH-sensitive transcriptional condensates, either by locating SEs directly linked to BRD4 condensates or by inferring target genes regulated through these sites. All approaches converged on a highly similar set of TRs, with RELA, IRF family, and STAT family consistently emerging as key transcriptional regulators. This convergence further supported by unsupervised clustering of chromatin regions, demonstrates that TRs identified by the region-centric analysis are reproducible and robust.

### RELA and IRF3 binding patterns within SEs align with BRD4 signals and target gene expression across pH conditions

To validate whether the identified TRs are involved in BRD4-associated, pH-sensitive transcriptional condensates, we examined RELA and IRF3 ChIP-seq data under LPS stimulation at pH 7.4 and pH 6.5. At pH 7.4, we identified 19,340 RELA binding sites in the genome, while under pH 6.5, we detected only 3,508 RELA peaks. For IRF3, we observed 8,191 peaks at pH 7.4 and only 3,752 peaks at pH 6.5, respectively. Both RELA and IRF3 showed markedly reduced binding under acidic conditions, indicating decreased activity at low pH. Centered on pH 7.4 peaks within pH-sensitive SE regions, we compared the signals across both pH 7.4 and pH 6.5 for the two transcription factors. The majority of peaks exhibited higher RELA and IRF3 occupancy at pH 7.4 than at pH 6.5 (Figures 4A, B), with most peak signals significantly reduced under pH 6.5 from pH 7.4 (Figures 4C, D).

**Figure 4.**
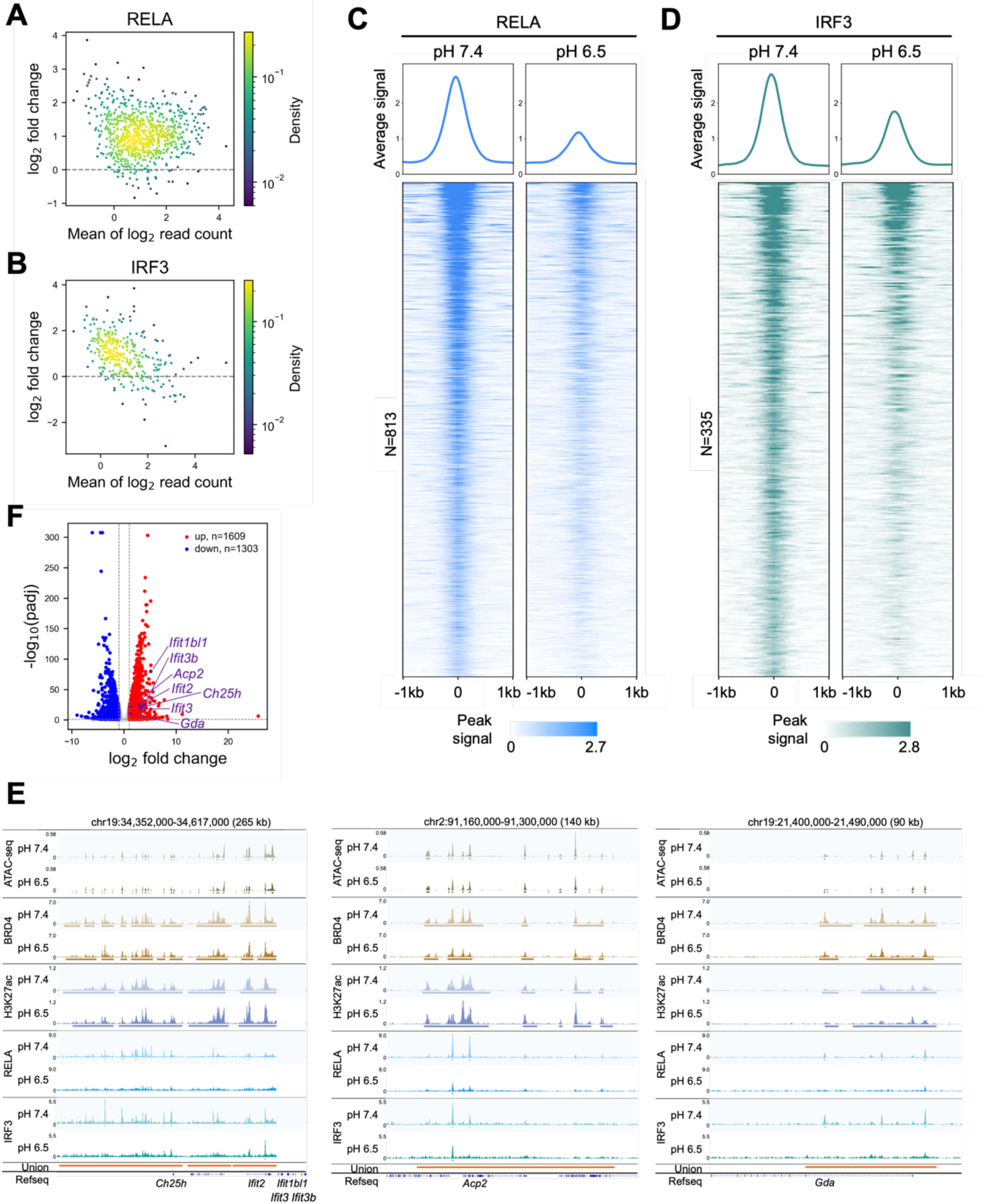
Validation of RELA and IRF3 as BRD4-associated, pH-sensitive transcriptional regulators. (**A**, **B**) MA plots of normalized ChIP-seq read counts comparing pH 7.4 and pH 6.5 at pH 7.4 peaks within pH-sensitive SE regions under LPS stimulation for Rela (**A**) and Irf3 (**B**). (**C**, **D**) Heatmaps of normalized ChIP-seq signals for Rela (**C**) and Irf3 (**D**) at pH 7.4 and pH 6.5, centered on pH 7.4 peak summits within pH-sensitive SE regions under LPS stimulation. (**E**) Examples of RELA and IRF3 ChIP-seq binding patterns at SE regions, aligning with BRD4 occupancy and corresponding changes in target gene expression across pH conditions. (**F**) Volcano plot of differentially expressed genes (DEGs) comparing pH 7.4 versus pH 6.5 under LPS stimulation. Genes marked in purple indicate significantly higher expression at pH 7.4, consistent with BRD4, Rela, and Irf3 binding changes shown in panel (**E**).

To examine how RELA and IRF3 binding correlates with potential target gene activation within BRD4-associated SE regions, we performed differential gene expression analysis for LPS-stimulated cells at pH 7.4 and pH 6.5. Several key immune-response genes, including *Ch25h*, *Acpt2*, *Gda*, and *Ifit* family, which are located within BRD4-associated, pH-sensitive SEs and defined as BRD4 condensate target genes (Figure 4E), had significantly higher expression level under pH 7.4 than pH 6.5 (Figure 4F). These loci also exhibited stronger RELA and IRF3 binding at pH 7.4, suggesting coordinated transcriptional regulation mediated by BRD4-associated, pH-sensitive transcriptional condensates.

In summary, we identified RELA and IRF3 exhibiting reduced binding in acidic environments and preferential localization within SEs. Their binding patterns align closely with BRD4 signals and target gene expression, providing functional validation that the identified TRs participate in transcriptional regulation mediated by BRD4-associated, pH-sensitive transcriptional condensates.

## Discussion

In this study, we systematically decoded TRs linked to BRD4-associated, pH-sensitive transcriptional condensates in macrophages through integrative chromatin analyses. Using multiple complementary analytical frameworks, we identified a convergent set of BRD4-associated, pH-sensitive TRs, including RELA, STAT family, and IRF family, under LPS stimulation. These factors are well known to play central roles in inflammatory signaling and immune regulation^14,19–22^, suggesting that BRD4-associated condensates act as regulatory hubs that integrate diverse transcriptional pathways under physiological conditions.

Using ChIP-seq, we validated that RELA and IRF3 show markedly reduced binding under acidic stress, consistent with decreased BRD4 occupancy at SE sites. This reduction corresponded to decreased expression of key immune-response target genes, including *Nfkb2*, *Tlr1*, *Tlr6,* and *CCL* family members. Together, these results provide functional evidence that RELA and IRF3 act as BRD4-associated, pH-sensitive TRs that coordinate gene activation through transcriptional condensates.

Recent studies^10,23^ have shown that BRD4 plays a key role in macrophage immune regulation by forming transcriptional condensates, which dissolve under acidic pH without affecting histone acetylation. In our systematic analysis, we identified numerous SE sites in the mouse genome that exhibit this pattern, which showing pH-sensitive BRD4 enrichment but stable H3K27ac levels, and the identified TRs were correspondingly enriched at these BRD4 pH-sensitive sites.

Overall, our study provides a comprehensive framework for identifying TRs functionally linked to BRD4-associated, pH-sensitive transcriptional condensates and demonstrates that environmental cues such as pH profoundly reshape condensate composition and transcriptional output. Although it remains unclear whether these TRs actively drive condensate formation or are recruited as functional components within existing condensates, our findings highlight a regulatory network through which BRD4 and associated TRs integrate environmental signals to control gene expression. These results provide important insights into how acidic stress modulates transcriptional condensates and macrophage immune responses.

## Materials and Methods

### Data processing

ATAC-seq (GSE269890), BRD4 ChIP-seq (GSE269299), H3K27ac/RELA/IRF3 ChIP-seq (GSE269891), and RNA-seq (GSE269298) datasets have been deposited to the NCBI Gene Expression Omnibus (GEO)^24^ with the corresponding accession numbers. All sequencing reads passed the Basic Statistics quality check using FastQC. All ChIP-seq reads were aligned to the mm10 reference genome using Bowtie2^25^ (Single-end: bowtie2 --very-sensitive -U SRR.fastq | samtools view -Sb -o SRR.bam; Paired-end: bowtie2 -x mm10 −1 R1.fastq −2 R2.fastq --very-sensitive --no-mixed --no-discordant | samtools view -Sb -o SRR.bam). Reads mapped to the mitochondrial DNA were discarded. Duplicate reads were removed using Picard^26^, and low-quality reads (MAPQ < 30) were filtered out using samtools^27^. Reads overlapping blacklist regions^28^ were also excluded.

For BRD4 and H3K27ac ChIP-seq data, broad peaks were identified using SICER2^11^ with two parameter sets: window size 1000 and gap size 3000 (sicer -t treatment_bed -c control_bed -s mm10 -w 1000 -g 3000); and window size 200 and gap size 600 for higher resolution.

For RELA and IRF3 ChIP-seq, narrow peaks were identified using MACS2^29^ (macs2 callpeak -f BAMPE -g mm -B --keep-dup 1 -q 0.01 --SPMR -t SRR_clean.bam). Bigwig files were generated using BEDTools^30^, deepTools^31^, and UCSC tools^32^.

For ATAC-seq, raw reads were first trimmed using Trim Galore (trim_galore --paired --nextera), Trimmed reads were aligned to mm10 using Bowtie2 (bowtie2 -x mm10 −1 R1.fq.gz −2 R2.fq.gz --very-sensitive -X 2000 --no-mixed --no-discordant | samtools view -Sb). After the alignment, reads mapped to chrM, duplicate reads, and low-quality reads (MAPQ < 30) were removed. Reads in blacklist regions were excluded. The filtered paired-end BAM files were converted to BEDPE format using MACS2 (macs2 randsample -i clean.bam -f BAMPE -p 100 -o clean.bed), Peaks were then called using MACS2 (macs2 callpeak -f BEDPE -g mm -B -q 0.01 --SPMR -t clean.bed).

For RNA-seq, reads were aligned to mm10 using STAR^33^ (STAR --runMode alignReads -- genomeDir mm10 --readFilesIn R1.fastq.gz R2.fastq.gz --outFilterMultimapNmax 1 -- outSAMtype BAM SortedByCoordinate --quantMode GeneCounts --readFilesCommand zcat), Gene expression levels were then quantified using RSEM^34^ (rsem-calculate-expression -- fragment-length-ma× 1000 --estimate-rspd --paired-end --bam out.bam rsem_reference).

Differentially expressed genes between pH 7.4 and pH 6.5 under LPS stimulation were identified using DESeq2^35^ with a threshold of fold change > 2 and adjusted P-value < 0.05.

### Region-centric analysis

Each ATAC-seq, BRD4, and H3K27ac dataset included four conditions representing combinations of LPS stimulation and pH (LPS-stimulated or control, at pH 7.4 or pH 6.5). Peaks for all factors from all conditions were merged to a single union peak set. Candidate super-enhancer regions were defined as the union peaks of at least 10 kb in length.

To identify BRD4-associated, pH-dependent condensate sites, we focused on regions showing significantly higher BRD4 signal under normal pH (7.4) compared to acidic conditions (6.5). For each union region, read counts were quantified and normalized to the total number of reads mapped to all union peaks within each sample. Differential analysis was performed using the SICER2 differential function to identify sites with significantly increased BRD4 signal (FDR < 0.01, log_2_FC > 0.25) under pH 7.4 compared to pH 6.5, but not significantly higher H3K27ac signal (excluded all sites with H3K27ac FDR < 0.01). These regions were defined as pH-sensitive SE regions. Union sites showing no significant enrichment of either BRD4 or H3K27ac signals were defined as pH-insensitive SE regions.

Using BRD4-associated pH-sensitive and H3K27ac pH-insensitive candidate transcriptional condensate regions as input, with each site was assigned a score corresponding to the BRD4 LPS 7.4 vs. 6.5 –log_10_ (FDR) from SICER2, we then applied *BART* in region mode to identify TRs significantly associated with either the pH-sensitive or pH-insensitive SE regions (P-value < 0.01). By comparing results from both sets, we identified TRs that were shared pH-sensitive and pH-insensitive SE regions, as well as those uniquely enriched in the pH-sensitive SE regions.

To infer target genes of the pH-sensitive and pH-insensitive SE regions, regulatory potential (RP)^13^ scores were calculated for each gene. For each chromatin mark, the RP of gene *i* was defined as the sum of the chromatin mark signals weighted by their genomic distance from the transcription start site (TSS). Specifically, ChIP-seq signals in 200-bp bins within ±100 kb of the TSS were collected, and each bin’s signal was weighted by a sigmoid decay function to compute the total RP for gene *i*:

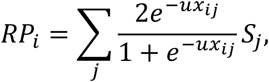

where *S*_j_ is the score of bin *j* surrounding the TSS of gene *i*, *x_*i*j_* is the genomic distance between the TSS and the center of bin *j*, and *u* controls the decay rate such that the half-life of the decay function is 10 kb.

Genes with non-zero RP for BRD4 from the pH-sensitive and pH-insensitive SE regions under the LPS pH 7.4 condition were considered target genes of the pH-sensitive and pH-insensitive SEs, respectively. Unique target genes were then identified for each set. Functional enrichment analysis was performed using DAVID^36,37^ to identify enriched gene sets and pathways.

Using the unique pH-sensitive and pH-insensitive SE regions target gene lists, we further applied BART^12^ in gene mode to identify TRs regulating these target genes (P-value < 0.01).

By comparing TRs identified from both analyses, we defined candidate regulators as those shared between the region-based and target gene-based analyses and uniquely enriched in the pH-sensitive SE regions, representing potential TRs associated with BRD4-associated, pH-sensitive transcriptional condensates.

### Gene-centric analysis

To identify genes potentially regulated by BRD4-associated pH-sensitive transcriptional condensates, we performed differential analysis based on the RP scores calculated in the region-centric analysis. Specifically, genes showing significantly higher RP scores under normal pH (7.4) compared to acidic pH (6.5) for BRD4 but not for H3K27ac were selected. Statistical significance was determined using a theoretical threshold derived from the MA plot, defined as:

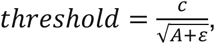

where c=1.5 and e=2 in this analysis. Genes exceeding this threshold were considered significantly enriched under normal pH. Selected genes were then analyzed using BART in gene mode to identify TRs significantly associated with BRD4-associated, pH-sensitive transcriptional condensate regulation.

### Open chromatin-centric analysis

Peaks from ATAC-seq across four conditions were merged to generate an ATAC-seq union peak set. For each union region, read counts were quantified and normalized to the total number of reads mapped to all union peaks within each sample. Because BRD4 and H3K27ac occupy broader regions, each ATAC-seq union site was extended by ±2 kb from its center. Read counts for BRD4 and H3K27ac were quantified within these extended regions and normalized by the total reads mapped to all extended sites.

Differential analysis was then performed to identify BRD4-associated pH-sensitive ATAC-seq union sites, defining pH-sensitive and pH-insensitive SE regions based on the theoretical threshold described in gene-centric analysis. BART was applied in region mode to identify TRs uniquely associated with the pH-sensitive SE regions.

To determine target genes of the pH-sensitive ATAC-seq union sites, RP scores were calculated for each gene using normalized read count on extended regions surrounding its TSS, following the same procedure as described in the region-centric analysis. BART in gene mode was subsequently used to identify TRs significantly associated with these pH-sensitive target genes. TRs identified by both the region-based and target gene-based analyses were considered candidate TRs associated with BRD4-dependent, pH-sensitive transcriptional condensates at open chromatin regions.

### Unsupervised clustering analysis

For each candidate super-enhancer site, a union profile was generated. This profile included the log_2_ fold changes of each chromatin mark between LPS-stimulated versus control conditions at fixed pH 7.4 or 6.5, and between pH 7.4 versus pH 6.5 at fixed LPS-stimulated or control conditions. These log_2_ fold-change values served as descriptive variables for each site. Unsupervised clustering was performed using k-means with k = 7 to group sites into seven clusters based on their chromatin mark profiles. For each site, odds ratios were calculated, and a one-sided Fisher’s exact test was used to assess whether pH-sensitive or pH-insensitive SE regions (as defined in the region-centric analysis) were significantly enriched in specific clusters. For ATAC-seq union sites within SE sites in each cluster, BART in region mode was applied to identify enriched TRs. The resulting TRs were then compared with those identified from the region-centric analysis.

### TR motif sites clustering analysis in the mouse genome

Motif information for TRs was collected from the JASPAR2024 CORE vertebrate non-redundant database (v2)^38^. FIMO^39^ was used to identify motif sites in the mm10 mouse genome with a threshold of --thresh 1e-4. Motif sites located within mouse genome blacklist regions were removed. BEDTools was used to identify the nearest downstream motif site for each motif, and the genomic distance between neighbor sites was calculated. Cluster Propensity (CP) score^8^ was computed to evaluate the genome-wide clustering tendency of motif sites, following our previously study^8^.

## Acknowledgements

The authors thank members of the Zang Lab and the Zhou Lab for inputs and comments to the manuscript. This work was partially supported by NIH grants R35GM133712 (C.Z.), R35GM151000 (X.Z.), state funding within the University of Virginia Comprehensive Cancer Center (C.Z.), Smith Family Foundation Odyssey award (X.Z.), and Boston Children’s Hospital Research Executive award (X.Z.).

## Supplementary Figures

**Supplementary Figure 1.**
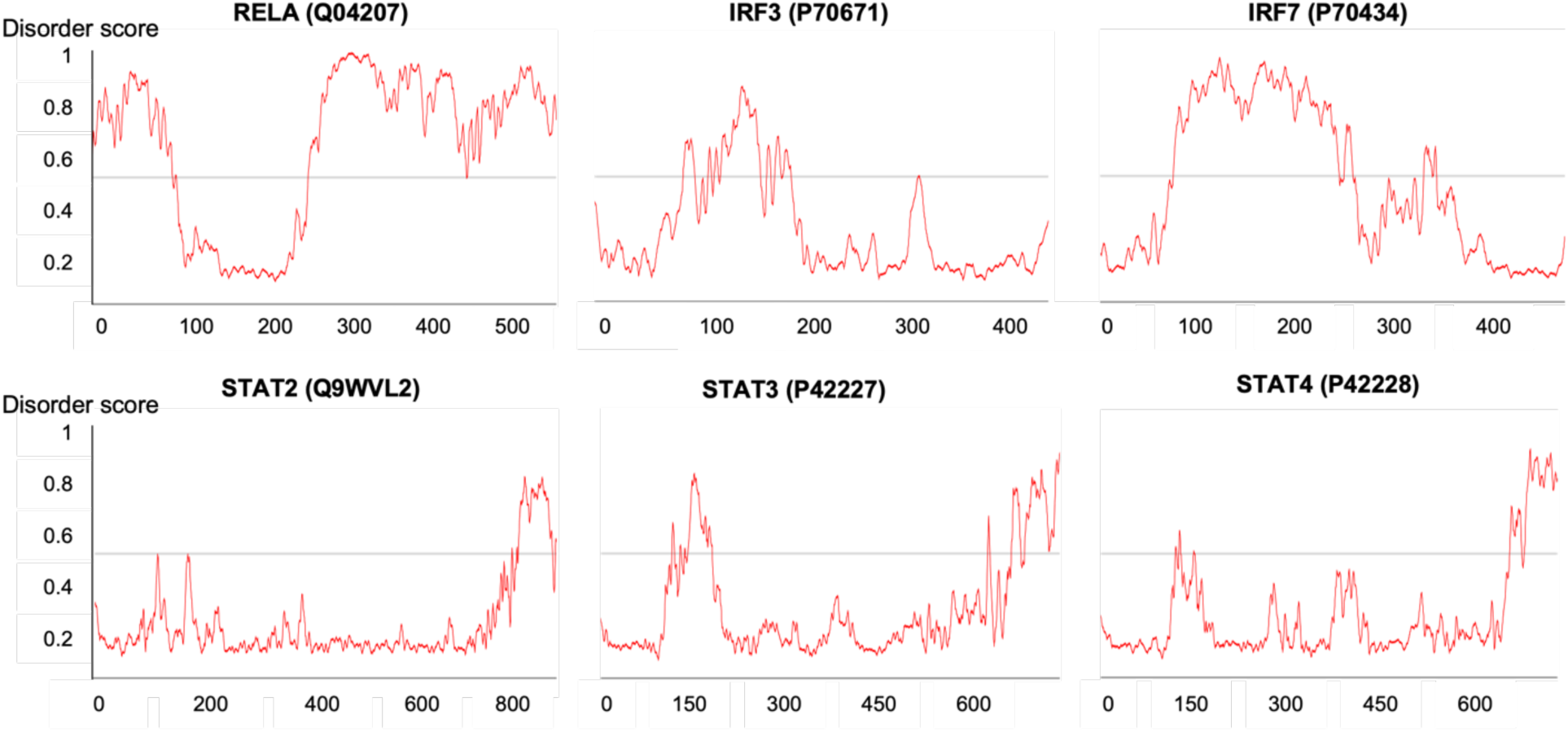
Prediction of intrinsically disordered regions (IDRs) in TF proteins using AIUPred. Regions with disorder scores above 0.5 are considered intrinsically disordered.

**Supplementary Figure 2.**
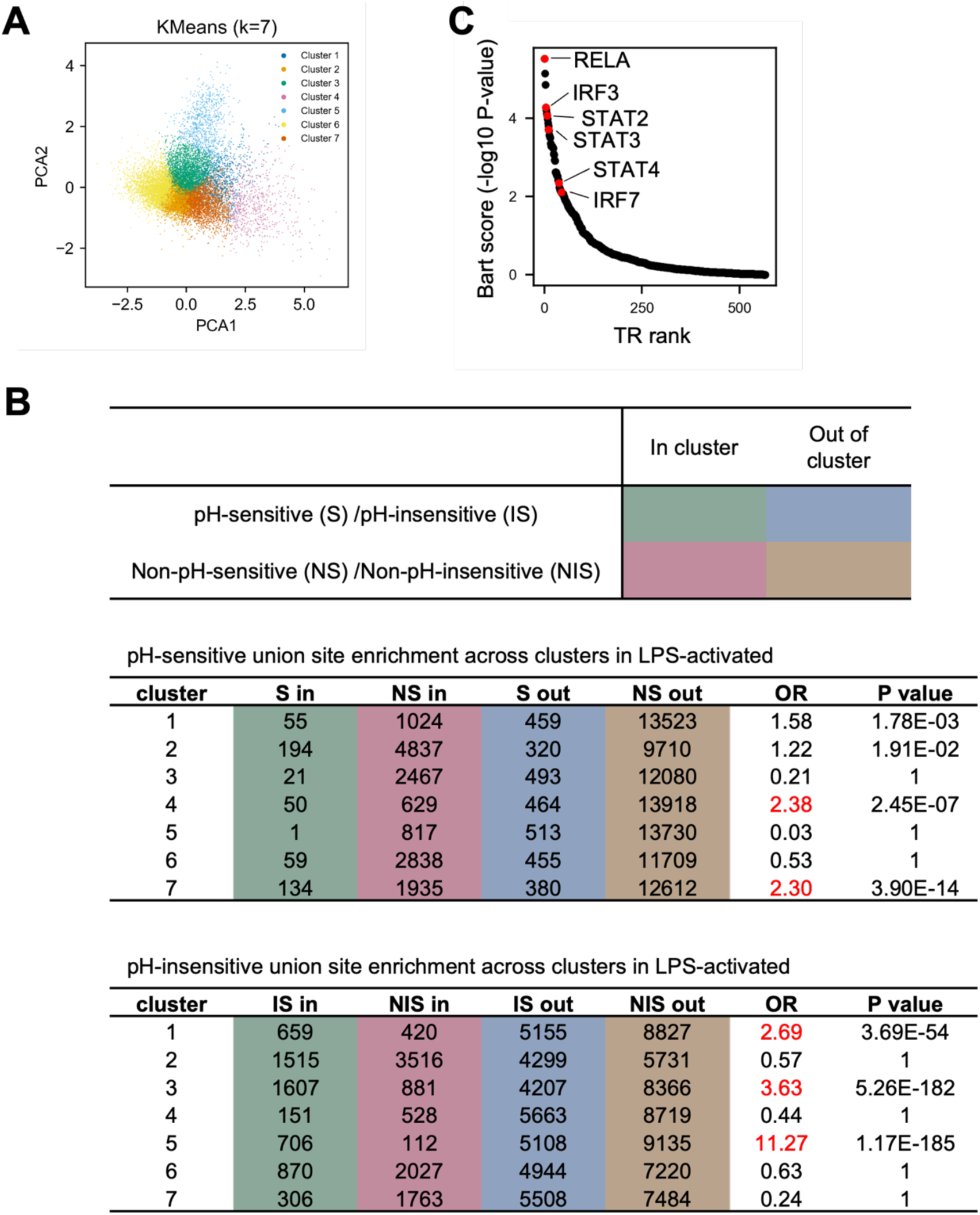
(**A**) *K*-means clustering of union site profiles. Scatter plot shows the clustering results projected onto the first two principal components (PCA1 and PCA2). (**B**) Enrichment analysis of *pH*-sensitive and *pH*-insensitive union sites identified in the peak-centric analysis within each *K*-means cluster. Clusters enriched for *pH*-sensitive or *pH*-insensitive sites with an odds ratio (OR) ≥ 2 are highlighted in red. (**C**) TRs enriched in cluster 7 ATAC-seq union peaks, with highlighted factors shared with the region-centric analysis.

